# Extensive recoding of dengue virus type 2 specifically reduces replication in primate cells without gain-of-function in *Aedes aegypti* mosquitoes

**DOI:** 10.1101/365189

**Authors:** Charles B. Stauft, Sam H. Shen, Yutong Song, Oleksandr Gorbatsevych, Emmanuel Asare, Bruce Futcher, Steffen Mueller, Anne Payne, Matthew Brecher, Laura Kramer, Eckard Wimmer

## Abstract

Dengue virus (DENV), an arthropod-borne (“arbovirus”) virus causing a range of human maladies ranging from self-limiting dengue fever to the life-threatening dengue shock syndrome, proliferates well in two different *taxa* of the Animal Kingdom, mosquitoes and primates. Unexpectedly, mosquitoes and primates have distinct preferences when expressing their genes by translation, e.g. members of these taxa show taxonomic group-specific intolerance to certain codon pairs. This is called “codon pair bias”. By necessity, arboviruses evolved to delicately balance this fundamental difference in their ORFs. Using the mosquito-borne human pathogen DENV we have undone the evolutionarily conserved genomic balance in its ORF sequence and specifically shifted the encoding preference away from primates. However, this recoding of DENV raised concerns of ‘gain-of-function,’ namely whether recoding could inadvertently increase fitness for replication in the arthropod vector. Using mosquito cell cultures and two strains of *Aedes aegypti* we did not observe any increase in fitness in DENV2 variants codon pair deoptimized for humans. This ability to disrupt and control an arbovirus’s host preference has great promise towards developing the next generation of synthetic vaccines not only for DENV but for other emerging arboviral pathogens such as chikungunya virus and Zika virus.

## Introduction

Synthetic biology has the potential to revolutionize the rapid development of vaccines to treat infectious diseases as the research paradigm shifts from empirical to rational design [1,2]. Since the first demonstration of an infectious fully synthetic virus in 2002 (7.5 kb) [3] and the ensuing initial societal concerns[4], rapid advances in DNA synthesis including decreased cost [5], has led to the general acceptance of synthetic organisms as a research tool[1,6,7]. This has also resulted in the rapid development of a new generation of synthetic vaccine candidates beneficial for humans and domesticated animals[1]. The conception of novel beneficial variants of existing organisms for the treatment of cancer[8,9] or for use as vaccines[10,11] have become new exciting branches in molecular medical research. However, there have been apprehensions over the use of this technology to synthesize dangerous pathogens by chemical synthesis such as poliovirus [4] or 1918 Influenza virus [12]. Alarm has been expressed about the ethics of using this technology: while there is great promise for the production of novel vaccines to improve human health there are also risks if the technology is misused, a dilemma referred to as “Dual Use Research”[13].

DENV is an enveloped, plus stranded RNA arbovirus (genome ~11 kb) of the genus *Flavivirus*, family *Flaviviridae*. DENV is separated into 4 serotypes and is primarily transmitted by the urban-adapted *Aedes aegypti* mosquito, a vector that has become widely distributed in tropical and subtropical regions. Efforts leading to an effective DENV vaccine have been complicated by the requirement that it must be tetravalent. Vaccination with a tetravalent vaccine can fortuitously lead to preferential protection against one serotype If followed by infection with a another single serotype, this may lead to severe or lethal disease mediated by antibody-dependent enhancement [14]. Complications with tetravalent DENV vaccines that have been recently reported call for new approaches to avoid unwanted outcomes [15,16]. Here we report further characterization of the first synthetic wild-type DENV2 based on strain 16681[17], and several *in silico* designed attenuated DENV2 variants carrying large-scale, but selective, genomic recoding of the open reading frame (ORF).

There are multiple methods of recoding a viral genome to achieve attenuation including the introduction of random point mutations [18], scrambling of codons while maintaining natural biases [19], reduction of codon bias for the host organism [20], and, as described here, changing of codon <u>**pair**</u> bias (CPB) to negative values[2,21]. Previously, our laboratory has exploited the universal phenomenon of CPB [22,23], whereby codons are prone to **pair** more or less frequently than expected with one another, independently of individual codon bias. Adjacent codons can form up to 36 different pairs that can encode the same pair of amino acids. The relative frequency of these pairs of codons can be represented by the natural logarithm of the ratio of the observed codon pair frequency to the expected codon pair frequency. This ratio is referred to as a codon pair score (CPS), and codon pairs that pair more frequently will have a positive “favorable” CPS while those unlikely to form a pair will have a more negative “disfavored” score. The nonrandom distribution of preferences for codon pairs is referred to as CPB [21]. Codon pair deoptimization (encoding an ORF largely with codon pairs with negative scores, see below), e.g. lowering the CPS, of a pathogen’s genome always results in attenuation across viral orders [17,24–29].

Available evidence suggests that CPB exists in all known taxa, including bacteria and yeast [22]. CPB for mammals is distinct from CPB in insects [17]. Arboviruses such as DENV, Zika virus, and chikungunya virus, must balance their CPB if they wish to replicate well in these different taxa. These organisms are optimal for studying the effects of altered CPB as they must contend with the translational machinery of primates and mosquitoes. It is possible to adjust the CPS of a virus to be negative (disfavored) with respect to mammals but be neutral with respect to insects. Previously, we described the *in vivo* and *in vitro* effects of reducing the CPS for DENV relative to the human genome while maintaining a neutral CPS for the *Aedes* genome [17].

Codon pair deoptimization (CPD) describes our method to introduce into a reading frame of the DENV coding sequence a large number of synonymous, disfavored (bad) codon pairs with negative codon pair scores without introducing mutations in the polyprotein or changing the use of existing codons. We note that the biological difference between “good” and “bad” codon pairs is small. However, if a large number of “bad” codon pairs are introduced into the ORF, the effect is disastrous for viral gene expression [21]. Our recoding strategy has raised a concern that dramatic recoding of the DENV genome may have unforeseen and dangerous consequences *vis a vis* the invertebrate vector. Whether through interaction between the RNAi based immune system of the mosquito [30] or some other means yet unidentified, there is a possibility that a sequence unknown to nature could have advantages that would enhance infection. We have, therefore, studied DENV variants for increased replication in mosquito cells. No apparent “gain-of-function” of these engineered strains was observed *in vitro* or *in vivo* using two different strains of *Ae. aegypti*.

## Results

### Synthesis and analysis of synthetic dengue type 2 infectious cDNA (DENV2^syn^)

Following *in silico* design, we synthesized three live-attenuated DENV2 viruses, each individually with *human* codon pair deoptimized coding sequences in one specific coding region of the polyprotein, yet leaving the insect-specific codon pair bias nearly wild type (Fig. 1). We refer to these DENV variants as E^hmin^, NS3^hmin^, and NS5^hmin^. All three proteins targeted here play multiple roles in the replicative cycle of DENV [31]. The number of nucleotide changes in, and the codon pair scores of, the recoded sequences E^hmin^, NS3^hmin^, and NS5^hmin^ are summarized in Table 1. We note that because only synonymous codons were moved codon bias (“codon usage”) remained unchanged. Similarly, the amino acid sequences of E^hmin^, NS3^hmin^, and NS5^hmin^ remained wild type. By design, the codon pair scores of the *hmin* constructs for all three proteins are significantly more negative than that of the wt sequences whereas the scores for insect ORFs remained almost wild type[17].

**Fig. 1.**
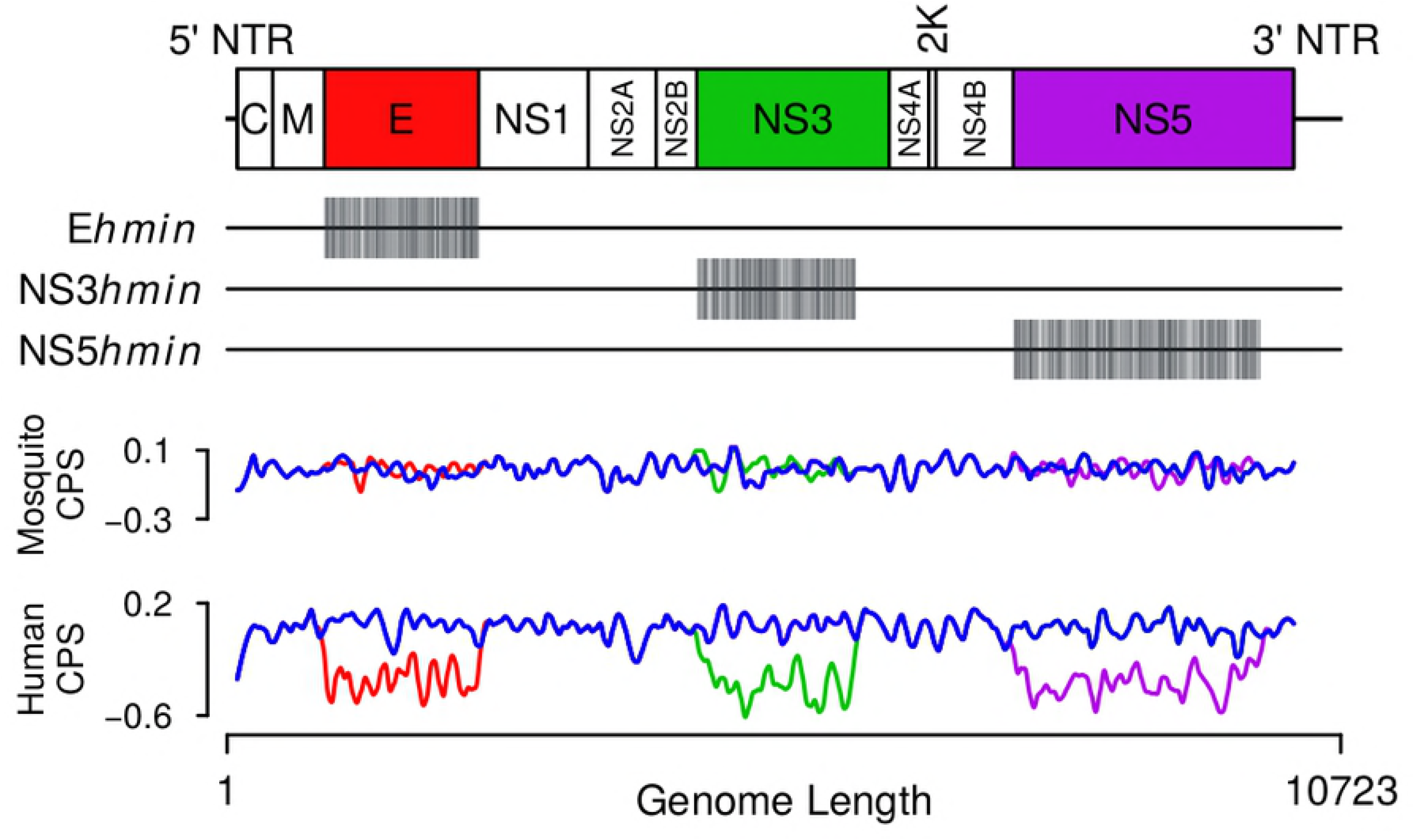
Diagram of the DENV2 genome marking the polyprotein coding region and the principle coding regions with the color-coded regions indicating the recoded regions.

**Table 1:**
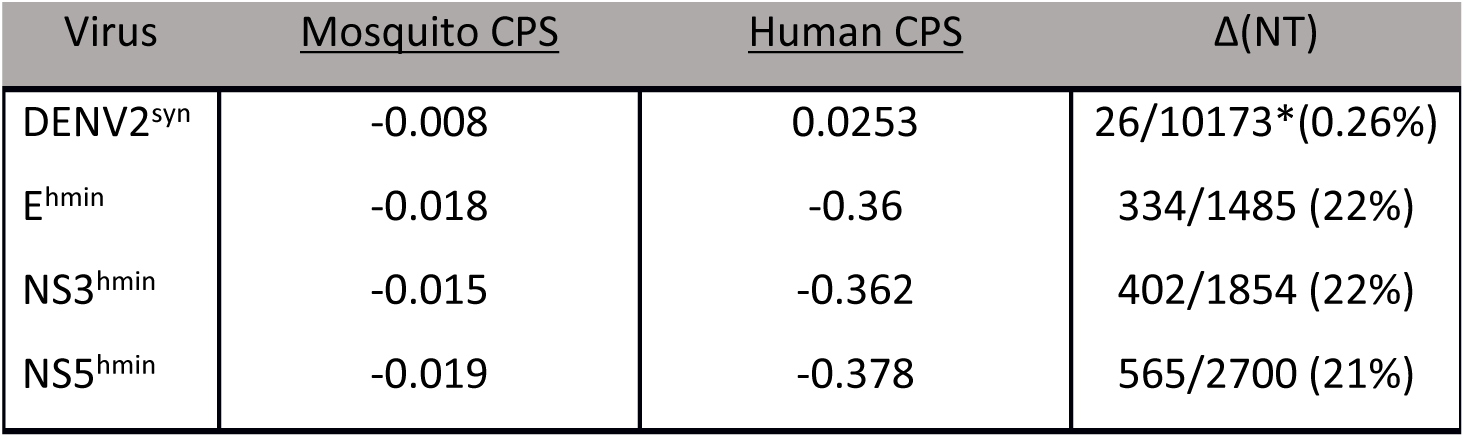
Relative CPS of each synthetic virus and mutations compared to wild-type. CPS is the natural logarithm of the ratio of the observed codon pair frequency to the expected codon pair frequency for the targeted region.

| Virus | Mosquito CPS | Human CPS | $\Delta$(NT) |
| --- | --- | --- | --- |
| DENV2<sup>syn</sup> | -0.008 | 0.0253 | 26/10173*(0.26%) |
| E<sup>hmin</sup> | -0.018 | -0.36 | 334/1485 (22%) |
| NS3<sup>hmin</sup> | -0.015 | -0.362 | 402/1854 (22%) |
| NS5<sup>hmin</sup> | -0.019 | -0.378 | 565/2700 (21%) |

### Replication of DENV2 E^hmin^, NS3^hmin^, and NS5^hmin^ variants in human macrophages and other primate cells

Macrophages are the predominant target cells in human dengue virus infections [32,33] and, as such, represent an *in vitro* model for attenuation of our synthetic DENV2 variants in the human host. Therefore, we focused on the growth properties of codon-pair deoptimized variants E^hmin^, NS3^hmin^, and NS5^hmin^ of DENV2^syn^ in cultured human macrophage cells as well as other cells of primate/human origin (Fig. 2). As our viruses were selectively deoptimized for attenuation in humans, we tested human THP-1 (Fig. 2A) and U937 (Fig. 2B) cells (induced monocyte-derived macrophages), the human lung epithelial cells A549 (Fig. 2C)[34], and two primate cells lines, Vero (Fig. 2D) and LLC-MK2 (Fig. 2E), for virus replication. Significant 1-2 logi_0_ reduction in Vero cells was observed with E^hmin^. Attenuation was far more pronounced for NS3^hmin^ and NS5^hmin^ (−5 log_10_ reduction in titer). With NS3^hmin^ infection of THP-1 cells, viral titers were static in cell culture supernatant, however, infection was confirmed using immune staining and visualization as shown by microscopy (Fig. S1). Together, these data confirm our hypothesis that CPD specific for human ORFs reduced the ability of the variants to proliferate in human and primate cells.

**Fig. 2.**
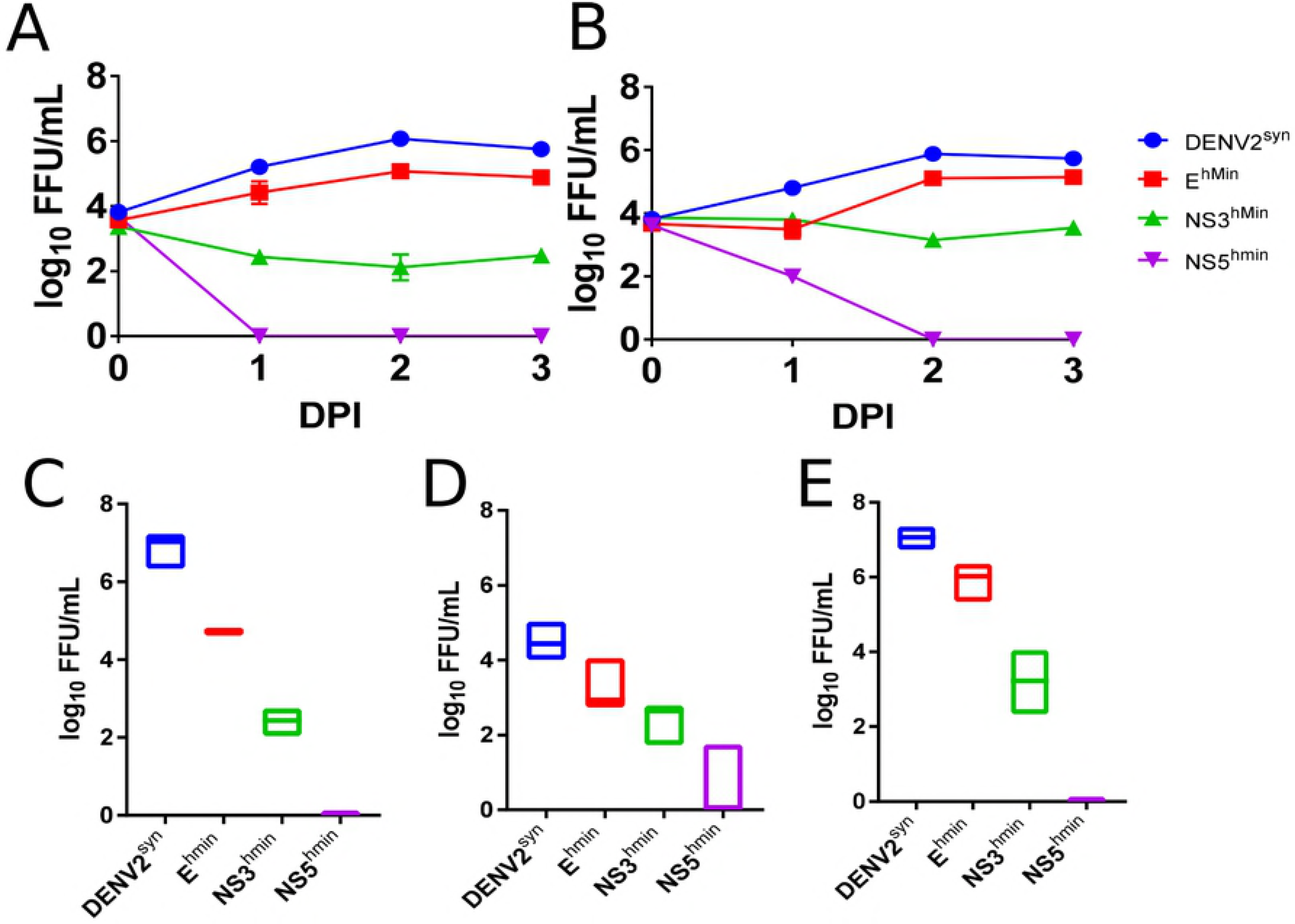
Replication of DENV2^syn^, E^hmin^, and NS3^hmin^ in human and primate cell lines. Human induced macrophage cell lines THP-1 (A) and U937 (B) were infected with each virus at a MOI of 1.0 and supernatant collected on 0-3 dpi (Days Post Infection) and titrated by FFA in C6/36 cells. Primate Vero (C) and LLC-MK2(D) as well as human lung epithelial A549 (E) cell lines were infected with each virus at a MOI of 1.0 and supernatant from 2dpi titrated by FFA in C6/36 cells.

### Replication of E^hmin^, NS3^hmin^, and NS5^hmin^ variants in mosquito cell cultures was not affected by our recoding

CPD leads to DENV2 variants with genetic sequences never before seen in nature [17]. Using a variety of human pathogens we have shown that human codon pair deoptimization for these agents leads to deficiencies in virus proliferation[35]. These DENV ORFs were deoptimized for human CPS, but, by design the CPS for *Ae aegypti* was left essentially wild-type. Thus, we do not expect this recoding to affect growth in mosquito-derived cells (15).

To investigate of selective attenuation by codon pair deoptimization, we originally used the mosquito cell line C6/36. This was derived from *Aedes albopictus* larvae and is now known to have mutations in the RNAi response. It has been proposed that these mutations (i.e., a lack of effective RNAi) facilitate infection by arboviruses [36]. Therefore, we carried out and report here studies using other mosquito cell cultures, specifically TRA-171 and CCL-125 cells. TRA-171 cells have been derived from the larvae of the non-blood feeding species of mosquito *Toxorhynchites amboinensis [37];* they have no identified deficiencies in insect innate immunity. CCL-125 cells have been derived from larvae of *Aedes aegypti* [38] and were initially reported to be refractory to DENV infection [39]. Recently, however, CCL125 have been shown to support DENV2 16681 replication after high MOI infection [40]. Our synthetic DENV2^syn^, which is based on DENV2 16681[17], also replicated to high (≤10^7^ FFU) titers in CCL-125 cells (Fig. 3B). Growth kinetics and virus production of DENV^syn^ and its human-deoptimized variants (E^hmin^, NS3^hmin^, and NS5hmin) were identical in TRA-171 (Fig. 3C), CCL-125 (Fig. 3D), and *Ae. albopictus* C6/36 cells (Fig. 3E) cells. This result strongly supports our hypothesis that human specific deoptimization indeed results in selective attenuation. We conclude that growth was poor in primate cells, wild-type growth is retained in mosquito cells (Table1).

**Fig. 3.**
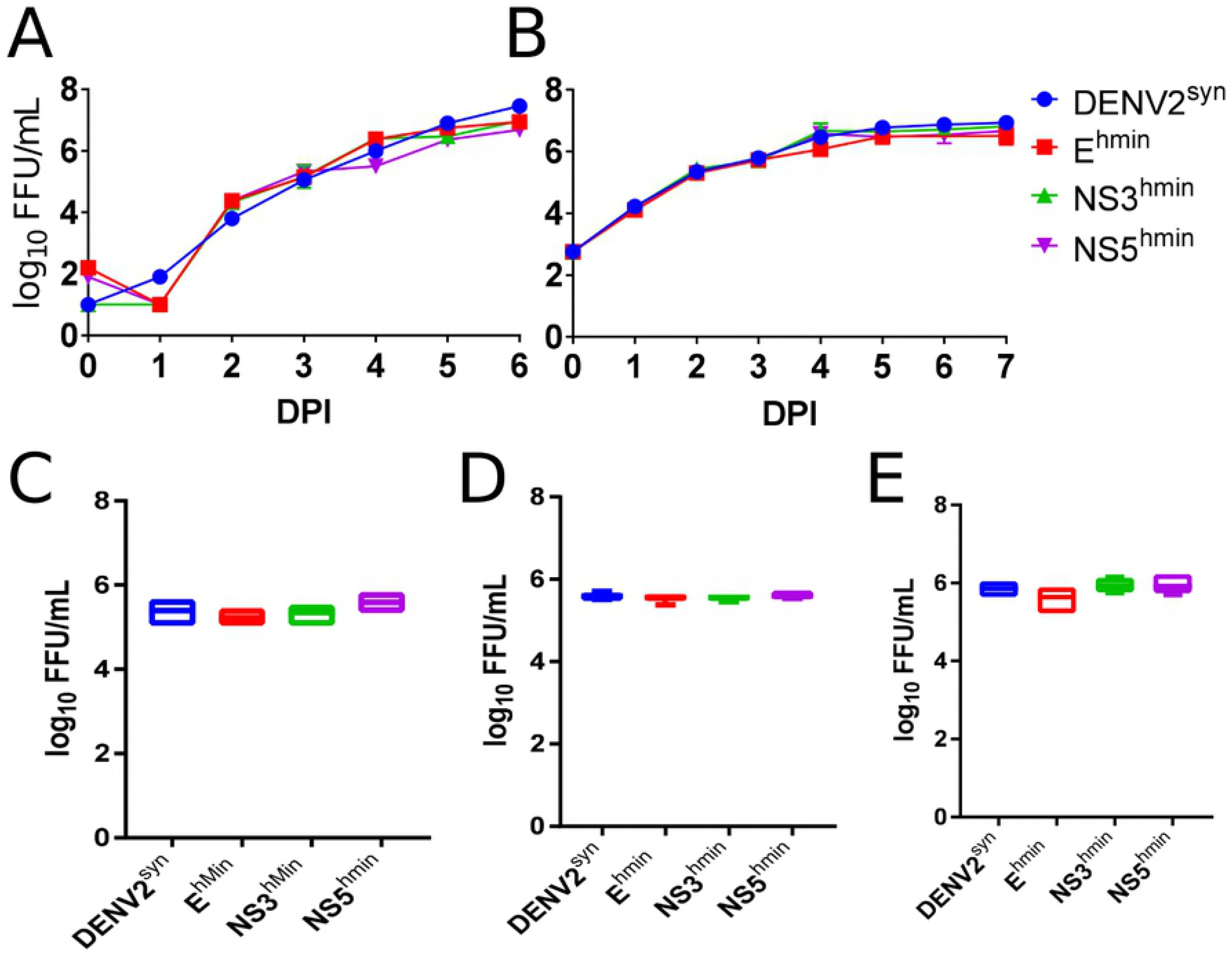
Growth kinetics of DENV2^syn^, E^hmin^, NS3^hmin^, and NS5^hmin^ in mosquito cell lines. Mosquito cell lines TRA-171 (A) and CCL-125 (B) were infected with each virus at a MOI of 0.1 or 1.0, respectively, and supernatant samples titrated by FFA daily for one-week post-infection. Mosquito cell lines TRA-171 (C), CCL-125 (D), and C6/36 (E) were infected with each virus at a MOI of 1.0, 5.0, or 1.0, respectively, and samples collected after incubation at 28°C for 2 days to compare titer by FFA in C6/36 cells.

### Infection and replication in Aedes aegypti mosquitoes were not affected by our recoding

To extend the results obtained in mosquito cell cultures, we also tested DENV2^syn^ and the human CPD variants in live *Ae. aegypti*. We tested the variants for changes in proliferation in two strains of *Ae. aegypti* mosquitoes using intrathoracic inoculation and infection *per os*. Viral stocks were diluted ten-fold from 200 to 0.2 FFU/ml in mosquito diluent solution and 4 μL inoculated into female *Ae. aegypti* (for more details see Materials and Methods). Mosquitoes were collected on day 5 and analyzed by Taqman RT-PCR to calculate the infectious-dose 50% (ID_50_). No significant difference in ID_50_ was observed between the DENV2^hmin^ variants and wild-type DENV2 16681 by t-test (P>0.09; Table 2).

**Table 2:** ID_50_ of intrathoracically inoculated *Aedes aegypti*.

| Input Titer/mL | -1.0<br>200 FFU | -2.0<br>20 FFU | -3.0<br>2 FFU | -4.0<br>0.2 FFU |  |
| --- | --- | --- | --- | --- | --- |
| Virus | % Infected |  |  |  | ID50 (Log10 FFU) |
| DENV2 16681 | 100 | 35 | 5 | 0 | -1.834 |
| DENV2<sup>Syn</sup> | 100 | 20 | 5 | 5 | -1.677 |
| E<sup>hmin</sup> | 95 | 30 | 5 | 0 | -2.172 |
| NS3<sup>hmin</sup> | 100 | 55 | 20 | 0 | -1.745 |
| NS5<sup>hmin</sup> | 95 | 25 | 0 | 0 | -1.671 |

We further tested viral replication in mosquitoes by using both intrathoracic injection (to bypass the midgut infection and replication barriers) as well as infectious bloodmeal (to measure viral replication under more natural conditions). Adult *Ae. aegypti* were intracranially injected with 2 × 10^4^ FFU virus. Mosquitoes were collected on days 3, 6,9, and 12 p.i. and analyzed by Taqman RT-PCR to see if there were any differences in total viral RNA titer. Among infected mosquitoes, the mean viral RNA titers were statistically similar at 3, 6, 9, and 12 days post infection with significantly but slightly reduced titers on days 3 (E^hmin^) and 12 (E^hmin^, NS3^hmin^) post-inoculation (Fig. 4). *Ae. aegypti* were then fed infectious bloodmeals containing ~4.0 × 10^6^ FFU/mL and tested for infection by Taqman RT-PCR. The rates of infection were low, but similar for each deoptimized virus compared to wild-type (Table 3).

**Table 3:** ***Aedes aegypti* infected with synthetic wild-type and deoptimized viruses**

| Virus | Bloodmeal Titer | Positive (%)<br>14 DPI |
| --- | --- | --- |
| DENV2<sup>Syn</sup> | $4.6 \times 10^6$/mL | 3/30 (10%) |
| E<sup>hmin</sup> | $5.2 \times 10^6$/mL | 3/30 (10%) |
| NS3<sup>hmin</sup> | $3.4 \times 10^6$/mL | 2/30 (7%) |
| NS5<sup>hmin</sup> | $1.6 \times 10^6$/mL | 1/30 (3%) |

**Fig. 4:**
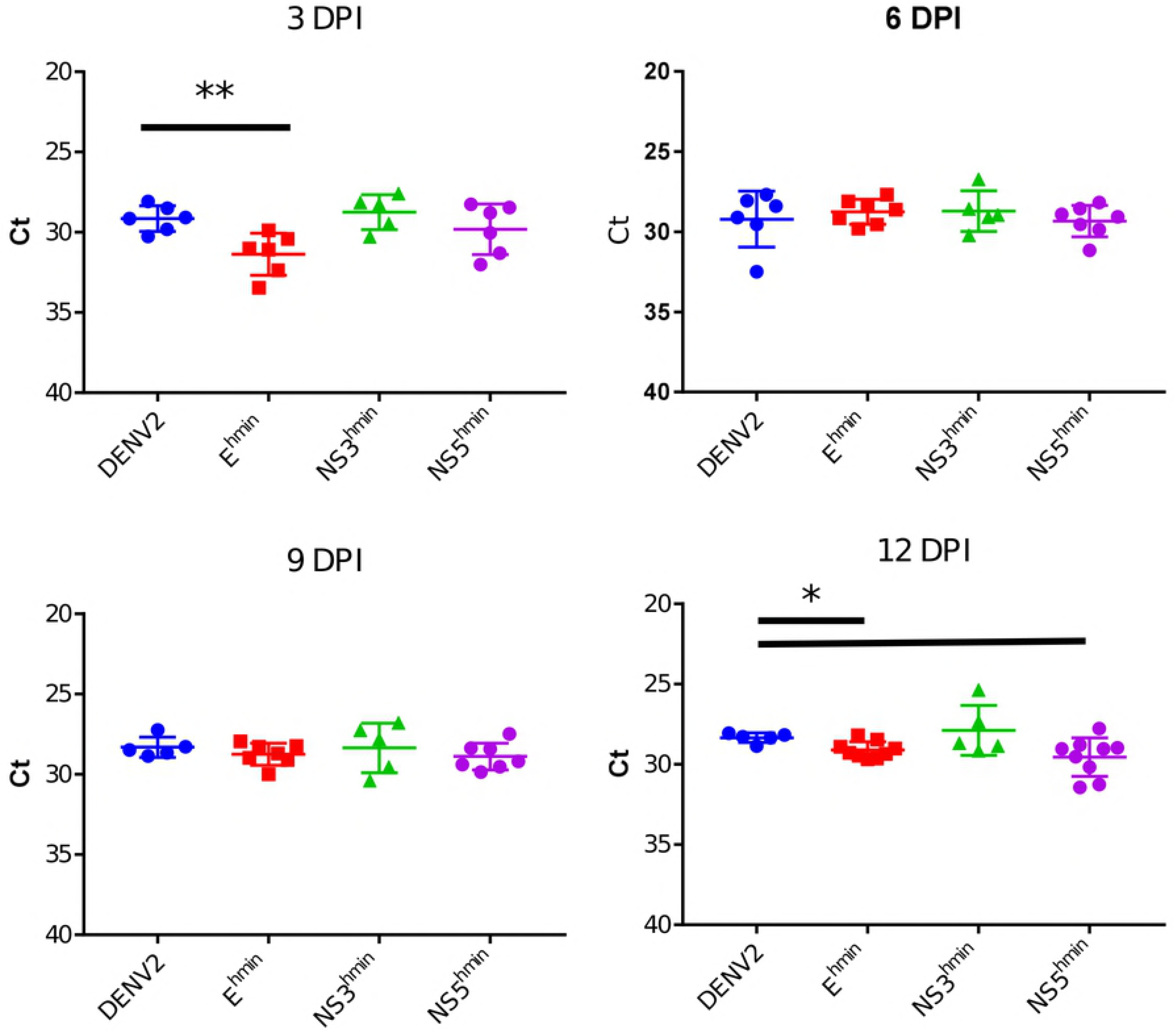
Infection kinetics in intrathoracically inoculated *Aedes aegypti*. Mosquitoes were inoculated with ~2 × 10^4^ FFU of each DENV strain and carcasses tested for viral RNA by Taqman qRT-PCR at 3, 6, 9, and 12 days post infection.

### Attenuation of CPD DENV by increased temperature sensitivity

The phenotype of temperature-sensitivity plays a significant role in some live virus vaccines as, for example, in live attenuated influenza vaccines [41] or the oral poliovirus vaccine [42]. Moreover, temperature sensitivity has been previously reported as a phenotype for codon-pair-deoptimized human respiratory syncytial viruses (RSV)[26,43]. Therefore, we examined deoptimized DENV2 variants E^hmin^ and NS3^hmin^ for temperature sensitivity in infected Vero cells (Fig. 5). As expected of selective human deoptimization, E^hmin^ and NS3^hmin^ replication was significantly reduced in Vero cells of primate origin (Fig. 5B). Because each variant replicates identically and forms equivalent FFU in C6/36 cells [17], all samples were assayed by FFA in C6/36 cells.

**Fig. 5.**
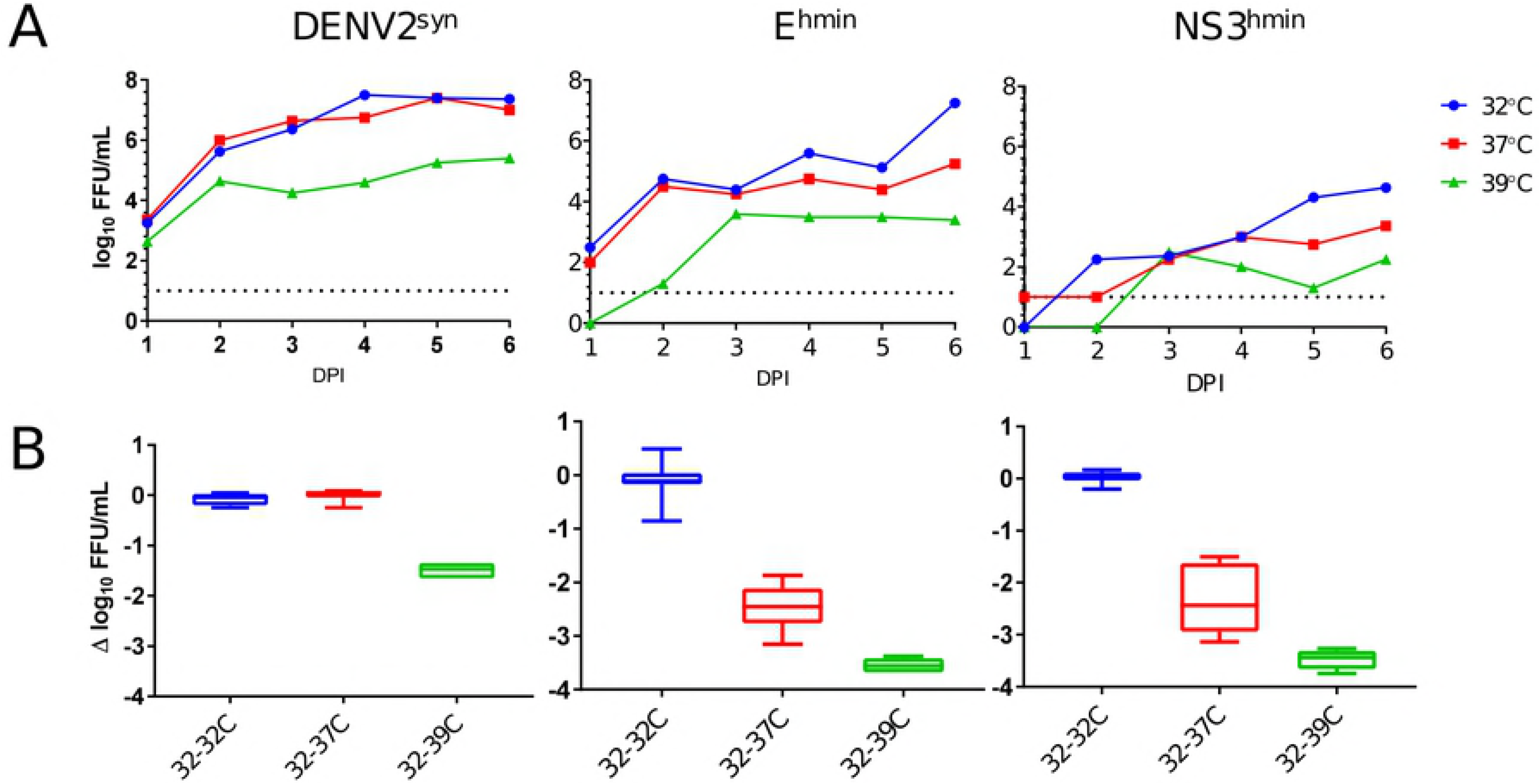
Growth curves at 32, 37, 39°C for DENV2^syn^, E^hmin^, NS3^hmin^ in Vero cells. Vero cells were infected at a MOI of 0.01 and infected cell supernatant titrated in C6/36 cells for 6 days post infection (A). Differences in titer between infected cells grown at 32°C and 37°C or 39°C were compared for each DENV variant (B).

Briefly, DENV2^syn^, E^hmin^, or NS3^hmin^ were used to infect Vero cells at a multiplicity-of-infection (MOI) of 0.01 at 32°C, 37°C, or 39°C (Fig. 5). Infected cells were incubated at the respective temperature and samples were taken daily for 6 days-post-infection (DPI). DENV2^syn^ replication was not affected by incubation at 32°C or 37°C (Fig. 5A), however, at 39°C a 2 log_10_ FFU/mL drop (Fig. 5B) in maximum titer was observed. In E^hmin^ infected cells, infectious virus was recovered at all three temperatures (Fig. 3A) but there was a 2-3 log_10_ FFU/mL drop between 32°C and 37°C and an even greater drop of nearly 4 log_10_ maximum titer (6 dpi) was observed between 32°C and 39°C (Fig. 3B). Infection of Vero cells with NS3^hmin^ revealed a similar trend in growth kinetics with reductions in maximum titer like those observed for E^hmin^ (Fig. 5). Temperature sensitivity experiments were attempted in C6/36 cells, however, the cultures were nonviable at 33°C and 37°C.

The spread of DENV2 variants in cell culture by focus forming assay (FFA) was also used to measure viral attenuation. FFA was conducted with each variant after simultaneous incubation at 28°C, 32°C, 37°C, or 39°C. Focus forming unit (FFU) size was not visibly different for DENV2^syn^ between 28°C and 37°C in Vero cells, however, a slight decrease in FFU size was observed at 39°C. This was in stark contrast for E^hmin^ FFU: at 32°C, 37°C, and 39°C the FFU size decreased dramatically with increasing temperature (not shown). FFU size at 28°C was like that at 32°C, and at no temperature was parity observed between DENV2^syn^ and E^hmin^ FFU size. The NS5^hmin^ and NS3^hmin^ variants did not produce FFU at any temperature after up to 8 days of incubation. Therefore, while DENV2 is somewhat temperature sensitive, the human deoptimized viruses appear to be more sensitive to increased temperature.

## Discussion

We have recoded individually three coding regions of the DENV2 polyprotein according to a master plan: increasing the number of “bad” codon pairs (negative codon pair scores) to reduce expression in primate cells while retaining a wild type average of codon pairs for wildtype expression in mosquito cells (Fig. 1A). Since wildtype DENV can grow well in *Ae*. species mosquitoes and humans, we speculated that the codon pairs used by DENV have a fine balance between invertebrate vector and vertebrate host. We have destroyed this calibrated, natural equilibrium through large-scale recoding (15) thereby generating DENV2 E^hmin^, NS3^hmin^, and NS5^hmin^ variants.

The calculated disruption of human codon pair score in CPD DENV2 variants resulted in attenuation using clinically relevant cultured human macrophage cells that we had not previously used. Moreover, CPD DENV2 variants were attenuated in a panel of different primate and human cell lines and are being tested for attenuation, immunogenicity and protection from challenge in Rhesus macaques at the Caribbean Primate Research Center in Puerto Rico (manuscript in preparation).

To address concerns about ‘gain-of-function’ and demonstrate the specificity of attenuation in CP-deoptimized DENV strains, we initially demonstrated in C6/36 cells, and have confirmed here in other mosquito cell lines as well as in two strains of live *Ae aegypti*, that our recoded DENV strains do not exhibit any unforeseen effects on replication. Our previous studies with DENV2^syn^ indicate that it is possible to leverage the codon pair bias difference between humans and mosquitoes to selectively attenuate the virus in one but not the other [17]. We hope to extend our studies to all four DENV serotypes and to other mosquito-borne viruses including emerging or re-emerging threats such as Zika virus [44], yellow fever virus, and chikungunya virus [45].

The mechanism of attenuation by codon-pair deoptimization is under investigation by our lab and others but remains to be solved. We suggest that it is likely the result of multiple factors; these include one or several parameters of the following:(i) the increased frequencies of CpG and UpA dinucleotides[46-48], possibly linked to an activation of the innate immune response, (ii) temperature sensitivity of the expression of recoded ORFs [26,29], (iii) stability of the recoded mRNAs[24,29], and (iv) possible complications with protein folding [28]. Unlike our previous work with Influenza A virus, dengue virus expresses its genome as a polyprotein. This is intriguing as the ribosome travels through the entire ORF thereby encountering individual segments of codon pair deoptimized sequences. Unlike the loss of protein expression observed in CPD Influenza viruses [24,29] or reduced protein expression in a codon pair deoptimized poliovirus variant [49]we do not have evidence of specific loss of recoded protein expression in DENV2^syn^ infected cells. Importantly, neither decreased temperature (Fig. 5), nor addition of innate immune inhibitor TBK-1 (not shown), nor addition of Jak Inhibitor I [17] completely rescued replication of the deoptimized DENV variants. However, in BHK (Baby Hamster Kidney) cells, which are deficient in RIG-I signaling [50], our viruses replicated identically to wild-type [17] which may indicate the role of this pathway in CPD DENV attenuation.

An intriguing report has recently been published that CpG dinucleotides are involved in the recognition of RNA by the Zinc-finger antiviral protein “ZAP”, leading to degradation of mRNA [18]. Indeed, in Table 4 we have enlisted the increase of CpG dinucleotide frequency in the CPD variants. The function of ZAP would link the innate immune response to the recognition of codon pair deoptimized viral genomes against DENV [18] and resulting degradation of viral RNA[57].

**Table 4:** Changes to CpG and UpA counts.

|  | DENV2<sup>syn</sup> | E<sup>hmin</sup> | NS3<sup>hmin</sup> | NS5<sup>hmin</sup> |
| --- | --- | --- | --- | --- |
| CpG | +0 | +67 | +84 | +98 |
| UpA | +0 | +31 | +48 | +49 |

Specifically relevant to DENV, CpG dinucleotides, while depleted in mammalian mRNAs, are observed with predicted frequency and show no downward bias in insects though UpA are depleted in the genomes of both insects and humans[58,59]. Although UpA frequencies were elevated in all three recoded DENV2 variants, there did not seem to be any adverse effects of this in either cell culture or live mosquitoes. These differences would impose contrasting selective pressures on DENV and other arboviruses which alternate replication in vertebrate hosts and arthropod vectors[60]. The CpG and UpA frequencies are more commonly found across codons in rare codon pairs, and so were increased in all of our CPD DENV2 variants. This increase was unavoidable and is very difficult to completely separate from changes due solely to codon pair bias, though there are current efforts in our lab to separate the two phenomena in poliovirus [49].

## Materials and Methods

### Construction of CPD and wild-type DENV2

Wild-type DENV2 virus was designed based on strain 16681 genome (Accession # U87411) and divided into four fragments (Fig. S1) incorporating 26 silent mutations (Table S1) as described previously [17]. CP-deoptimized viruses were recovered from transfection of C6/36 cells with capped RNA transcribed using HiScribe T7 *in vitro* transcription kit (New England Biolabs). Virus stocks were grown in C6/36 cells by harvesting cell culture supernatant 6 days post-infection at an MOI of 0.1. Viruses were titrated in C6/36 cells using a focus forming assay [17].

### Cell Cultures and Virus Production

Vero E6 (CCL-81), A549 (CCL-185), LLC-MK2 (CCL-7), and BHK-21 (CCL-10) cells were acquired from the American Type Tissue Culture collection (ATCC, Manassas VA) and grown in Modified Eagle’s Medium (MEM) supplemented with 10% fetal bovine serum (FBS; Gemclone) and Penicillin/Streptomycin (CellGro). U-937 (CRL-1593.2) and THP-1 (TIB-202) monocyte cells were grown in RPMI-1640 with 10% FBS and no antibiotics. U-937 and THP-1 cells were induced to become macrophages prior to experimentation as described previously [61]. All mammalian cell lines were maintained at 5% CO_2_ and 37°C. C6/36 cells (ATCC) were grown in Modified Eagle’s Medium (MEM) supplemented with 10% FBS, 1% non-essential amino acids (NEAA; Gibco), and 1x Penicillin/Streptomycin. *Toxorhynchites amboinensis* TRA-171 (CRL-1591) cells and *Ae. aegypti* CCL-125 cell lines were acquired from the ATCC. TRA-171 cells were grown in a 1:1 mix of L-15 and Mitsuhashi/Maramorosch medium (ATCC) supplemented with 2mM Glutamine,0.05% BSA, 1% NEAA, and 10% heat-inactivated FBS. CCL-125 cells were grown in MEM supplemented with 1% NEAA, Penicillin/Streptomycin, and 20% FBS. C6/36 and CCL-125 cells were incubated at 28°C and 5% CO_2_ while TRA-171 cells were incubated at 28°C without CO2. Growth kinetics were examined in each cell line by infecting the cells with each virus at a MOI of 1.0, collecting supernatant daily for 5-10 days, and titrating the virus in C6/36 cells using a focus forming assay (FFA).

### Infection of Aedes aegypti mosquitoes

*Ae. aegypti* mosquitoes were kindly provided by G.D. Ebel, Colorado State University, Fort Collins, CO, USA, originally collected in Poza Rica, Mexico. 3-5 day old females were fed infectious blood meals containing 4×10^6^ FFU/mL of DENV2^syn^, E^hmin^, NS3^hmin^, or NS5^hmin^. Blood meals were held at 37°C during feeding using a Hemotek Apparatus (Discovery Workshops). Engorged mosquitoes were sorted and maintained at 27°C with 16:8 hours light:dark cycle. Thirty individuals per group were collected at 5 days post infection for RNA isolation and detection using Taqman qRT-PCR. For the ID_50_ experiment, an established laboratory colony of *Ae. aegypti* was infected by intrathoracic injection with DENV2^syn^, E^hmin^, NS3^hmin^, or NS5^hmin^. Viral stocks were diluted 10-fold from 200 FFU/ml to 0.2 FFU/ml in mosquito diluent solution and 4 ul injected into each mosquito. Twenty surviving mosquitoes from each group were collected on day 5 p.i. and analyzed by Taqman PCR. A non-linear sigmoidal dose response curve was generated to calculate the ID_50_ for each virus which were compared using Student’s t-test (GraphPad Prism v 7.04, La Jolla, CA). Additionally, the Mexican strain of *Ae. aegypti* was infected by intrathoracic injection with ~2 × 10^4^ FFU of DENV2^syn^, E^hmin^, NS3^hmin^, or NS5^hmin^ as previously described [62]. Ten mosquitoes per group were collected on days 3, 6, 9, and 12 and analyzed by Taqman and FFA to see if there is any difference in rate of infection or total titer in the mosquitoes.

## Acknowledgements

We thank our colleagues J. Cello and J. Mugavero for discussion of aspects of this work.

